# Anterior cingulate cortex engram drives Post traumatic stress disorder-like memory impairments in a rodent model of traumatic fear

**DOI:** 10.64898/2026.09.01.748515

**Authors:** Flávia V. Simões, Lorenzo Baltimore, Zora Pelloquin-Mvogo, Laurent Groc, Aline Desmedt, Olivier Nicole, Sophie Tronel

## Abstract

Fear memories in post-traumatic stress disorder are marked by persistent cue-driven recollection and impaired contextual recall, yet the engram organization underlying this maladaptive state remains unclear. Here we used a rodent model combining contextual fear conditioning with systemic corticosterone to mimic trauma-associated glucocorticoid exposure. This paradigm generated a PTSD-like phenotype characterized by hypermnesia for a trauma-related but irrelevant (non-predictive of the threat) cue and contextual amnesia. Activity-dependent tagging and reactivation mapping revealed that traumatic memory is supported by a regionally dysregulated engram pattern, with enhanced recruitment of the anterior cingulate cortex (ACC) and basolateral amygdala (BLA), and reduced engagement of the dentate gyrus (DG). These changes persisted over time and correlated with the severity of the behavioral phenotype. Chemogenetic inhibition of ACC engram cells abolished traumatic memory expression, restored contextual recall, and normalized engram reactivation across DG and BLA. In contrast, inhibition of randomly tagged ACC or hippocampal populations had no such effect, indicating that the ACC engram is specifically required for traumatic memory expression. Together, these findings show that PTSD-like memory is not simply an amplified fear trace, but a distinct maladaptive engram state distributed across cortical-hippocampal-amygdalar circuits.

## INTRODUCTION

Emotional experiences are typically remembered more persistently than neutral ones (McGaugh, 2004; LaBar and Cabeza, 2006), an adaptive feature that enhances survival by guiding future behaviour away from potential threats (Tovote et al., 2015). However when emotional memory becomes excessive, it can become poorly contextualized, and thus maladaptive. This is particularly evident in post-traumatic stress disorder (PTSD), where trauma-related memories are repeatedly re-experienced in safe environments, while memory for the original contextual details remains fragmented or impaired (Brewin et al., 1996; Kaouane et al., 2012; Desmedt et al., 2015). PTSD can arise following severe stress exposure such as combat, sexual assault, or childhood abuse, and represents a major public health concern (Hoppen et al., 2024). Converging evidence from neuroimaging studies implicates dysfunction in brain circuits underlying PTSD-related memory, particularly within hippocampal– amygdala-prefrontal networks (Ben-Zion et al., 2023; Hinojosa et al., 2024).

At the cellular level, fear memories are believed to be encoded by sparse neuronal ensembles or engrams (Tonegawa et al., 2015; Josselyn and Tonegawa, 2020), whose reactivation is necessary and sufficient to trigger recall. While rodent studies have significantly advanced our understanding of these fear engrams, traumatic memories have been largely conceptualized as simply intensified versions of canonical fear memories. This perspective fails to explain a defining feature of PTSD: the paradoxical coexistence of heightened cue-driven recollection and impaired contextual memory, and thus the qualitative alteration of declarative fear memory (Kaouane et al., 2012; Al Abed et al., 2020). As a result, the organization of the neuronal ensembles supporting maladaptive traumatic memories remains poorly understood.

To address this gap, we used a rodent model that reproduces key features of PTSD-like memory by combining contextual fear conditioning with systemic corticosterone administration (Kaouane et al., 2012), thereby mimicking the glucocorticoid surge associated with trauma. We then characterized the engram organization underlying traumatic memory and showed that it fundamentally differs from that of adaptive fear memory. Specifically, traumatic memories recruit a regionally dysregulated engram pattern, marked by the preferential reactivation of the anterior cingulate cortex (ACC) and basolateral amygdala (BLA), together with disengagement of the dentate gyrus (DG) of the hippocampus. Furthermore, chemogenetic inhibition of ACC engram cells prevents the expression of traumatic memory and normalize engram, thereby promoting a normal adaptive fear memory profile. These findings reveal that PTSD-like memory is not merely an intensified fear trace but rather a distinct maladaptive engram state distributed across cortical-hippocampal-amygdalar circuits.

## RESULTS

### Corticosterone combined with contextual fear conditioning induces a PTSD-like memory characterized by enhanced BLA and ACC engram reactivation and reduced DG reactivation

Many studies investigating fear memory engram frequently equate generalized and exacerbated fear with PTSD-like syndrome, thereby often failing to capture the cardinal qualitative memory alteration observed in PTSD patients. To address this critical gap, we used a rodent behavioural paradigm previously established that effectively recapitulates both the contextual amnesia and the hypermnesia for trauma-related, but not predictive cue, characteristic of the disorder (Kaouane et al., 2012). This task adapts a standard contextual fear conditioning protocol wherein two auditory tones are presented but do not predict shock delivery; the tones serve to heighten the subject’s environmental salience without acquiring intrinsic aversive value, leaving the context as the sole valid predictor of the threat. Immediately after conditioning, mice received either an intraperitoneal injection of corticosterone (Cort) or Saline (Sal) together with 4-OHT for activity-dependent temporally-restricted neuronal tagging in TRAP2;Ai14 mice. (Fig. 1a,b). When exposed to the tone four days later in a neutral familiar context, Cort-injected mice exhibited an abnormal increased freezing level upon tone presentation, whereas Sal-injected controls showed no marked response to the tone (Fig. 1c). This effect was confirmed by calculating a tone ratio comparing freezing before and after tone presentation, which was significantly increased in Cort-injected mice (Fig. 1d) revealing a profound hypermnesic response to the non-predictive and thus irrelevant cue in the Cort cohort. Conversely, when re-exposed to the original conditioning context the following day, Cort-treated mice displayed a marked reduction in freezing across the 6-minute session compared to controls, indicative of contextual amnesia (Fig. 1e). This contextual deficit was particularly pronounced during the first two minutes of context exposure, where Cort-injected mice displayed significantly lower freezing levels (Fig. 1f). To evaluate the expression of this dual-phenotype across tests at the individual level, we calculated a “PTSD score” combining tone hyper-responsiveness and contextual memory impairment. This score was significantly increased in Cort-injected mice, demonstrating a robust and reliable shift toward a PTSD-like memory profile. (Fig. 1g). Both male and female mice were used in these experiments. The lack of sex-dependent differences demonstrates, for the first time, that female mice also exhibit a PTSD-like memory profile in this paradigm (Suppll Fig. 1a).

**Fig. 1.**
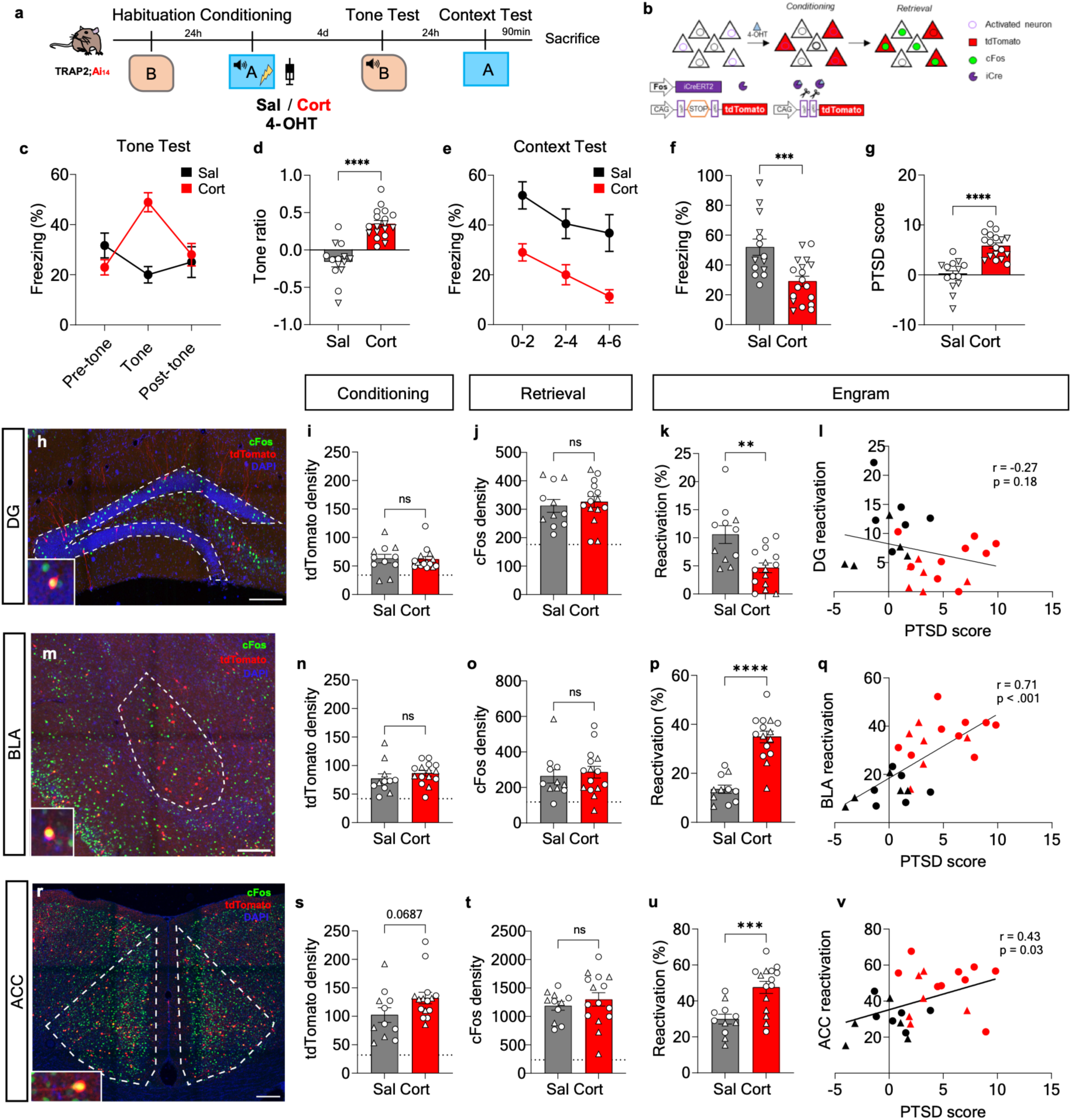
Traumatic fear induces hyper-reactivation of engram cells in the BLA and ACC alongside decreased reactivation in the DG. **a**, Experimental protocol for recent memory assessment. *Trap2;Ai14* mice were habituated to a neutral context (CtxB). The following day, mice were conditioned in CtxA and immediately received either Corticosterone (Cort, *n* = 16) or Saline (Sal, *n* = 11) injections combined with 4-OHT administration. Mice were tested 4 days later in CtxB in the presence of the tone, and evaluated the following day in the original conditioning context (CtxA). **b**, Schematic representation of activity-dependent recombination in *Trap2;Ai14* mice. **c**, Percentage of freezing in CtxB. Cort mice exhibited an elevated fear response during tone presentation compared to Sal mice *(***p* < 0.001). **d**, Tone ratio, which was significantly higher in Cort mice *(****p* < 0.0001). **e**, Percentage of freezing in CtxA. Cort mice displayed a decreased total freezing level compared to Sal mice *(***p* < 0.001, main effect of group). **f**, Freezing response during the first two minutes in CtxA, which was significantly lower in Cort mice *(***p* < 0.001). **g**, Calculated PTSD scores, showing a marked increase in Cort mice compared to Sal mice *(****p* < 0.0001). **h-t**, Histological assessment of engram and cellular activation across brain regions. Representative images (h, m, r; scale bar = 200 gm), quantified tdTomato÷ engram cell density (i, n, s), and c-Fos+ cell density (j, o, t) in the dentate gyrus (DG; h-j), basolateral amygdala (BLA; m-o), and anterior cingulate cortex (ACC; r-t). Densities did not significantly differ between groups (dashed lines represent baseline activation), k, I, Engram reactivation in the DG. DG tdTomato÷ cells showed significantly decreased reactivation in Cort mice during recall (p < 0.01) (k), which did not correlate with individual PTSD scores (I), p, q, u, v, Engram reactivation and behavioral correlations in the BLA (p, q) and ACC (u, v). Cort mice displayed significantly higher reactivation of tdTomato÷ cells compared to Sal mice, which positively correlated with individual PTSD scores (***p < 0.001, *\*\*\*\*p* < 0.0001). For histology and reactivation data: Cort, *n* = 15; Sal, *n* = 11. (For statistical details see table 1).

We then examined how this PTSD-like behavioural phenotype was associated with the recruitment and reactivation of memory engram cells. For this purpose, we used TRAP2;Ai14 mice, which allow temporally restricted, activity-dependent neuronal labelling. In this mouse line, CreERT2 expression is driven by the cFos promoter, and, in the presence of 4-OHT, activated neurons undergo Cre-mediated recombination leading to permanent tdTomato expression (Fig. 1b) (DeNardo et al., 2019). By administering 4-OHT immediately after conditioning, we labelled neurons recruited during memory encoding. Mice were then perfused 90 min after the context retrieval session, allowing us to quantify cFos expression as a marker of neuronal activation during recall. To control for possible side effects of 4-OHT, as it is an oestrogen receptor modulator, we injected both males and females with oil (vehicle) or 4-OHT, seeing no significant impact in PTSD score in either sex (Suppl Fig 1b).

We focused on brain regions classically implicated in fear memory and PTSD-related processes, including the dentate gyrus (DG), basolateral amygdala (BLA), and anterior cingulate cortex (ACC) (Fig. 1h,m,r). The density of tdTomato-positive cells did not significantly differ between Sal-and Cort-injected mice in the DG, BLA, or ACC, indicating that the overall size of the encoding-related neuronal population was not broadly altered by corticosterone treatment (Fig. 1i,n,s). To further assess whether these effects reflected regionally restricted changes in the density of tagged neurons, we quantified tdTomato-positive cell density across additional anatomical subdivisions. No major differences were detected between groups in the right or left BLA, right or left LA, dorsal or ventral DG, or posterior ACC (Suppl Fig. 1c-h,j). However, tdTomato density was increased in the anterior ACC, but not posterior, while mPFC showed a trend towards an increase (Suppl Fig. 1i,k) These complementary analyses indicate that the main PTSD-like phenotype is not explained by a global increase in tagged neuronal density across the fear memory network, although they suggest a preferential recruitment of anterior cortical areas after corticosterone treatment (Suppl Fig. 1). Similarly, total cFos density during recall was unchanged across these regions (Fig. 1j,o,t). However, the proportion of tdTomato-positive cells reactivated during recall was strongly region-dependent. In the DG, Cort-injected mice showed a significant reduction in engram reactivation compared with Sal controls (Fig. 1k). In contrast, Cort-injected mice exhibited a marked increase in engram reactivation within both the BLA and ACC (Fig. 1p,u). mPFC showed no difference in reactivation between groups (Suppl Fig. 1l,m). This reactivation pattern was largely consistent across sexes (Suppl Fig 1n-p) despite a baseline difference in the DG, where control (Sal) females exhibited lower reactivation levels than control males. To determine whether these region-specific changes in engram reactivation were related to the severity of PTSD-like memory impairment, we correlated reactivation levels with the individual PTSD-like score. DG reactivation did not significantly correlate with the PTSD-like score (Fig. 1l). By contrast, BLA and ACC reactivation showed positive correlations with the PTSD-like score, indicating that stronger reactivation of these regions was associated with more pronounced trauma-related hypermnesia and contextual memory impairment (Fig. 1q,v). Together, these findings suggest that corticosterone-induced PTSD-like memory is not associated with a generalized expansion of the fear memory engram, but rather with an abnormal redistribution of engram reactivation across the fear-memory network, characterized by reduced DG reactivation and enhanced BLA and ACC reactivation.

### Inverting recall session order does not alter PTSD-like memory expression or ACC engram reactivation

We next asked whether the PTSD-like memory phenotype depended on the order in which the two retrieval sessions were performed. To address this, we inverted the testing sequence by first exposing mice to the conditioning context A, followed 24 h later by tone presentation in the neutral familiar context B (Suppl Fig. 2a). Under this reversed testing order, Cort-injected mice still displayed reduced freezing upon re-exposure to the conditioning context compared with Sal-injected controls, indicating persistent contextual memory impairment (Suppl Fig. 2b,c). When tested the following day in the neutral context, Cort-injected mice also maintained an increased freezing response to the tone, as reflected by a significantly higher tone ratio (Suppl Fig. 2d,e). Accordingly, the integrated PTSD-like score remained significantly elevated in Cort-injected mice compared with Sal controls (Suppl Fig. 2f). These results indicate that the dissociation between hyper-responsiveness to the trauma-associated cue and impaired contextual memory does not depend on the order of recall sessions.

We then examined whether the engram alterations previously described were also preserved when the order of retrieval was inverted. cFos expression allows to inquire neuronal activation associated with tone recall. Under these conditions, Cort-injected mice showed increased reactivation of tdTomato-positive cells in the ACC compared with Sal controls, which was correlated with the PTSD score (Suppl Fig. 2 g,h). Thus, enhanced ACC engram reactivation is not restricted to one specific recall sequence and is also observed when the trauma-associated cue is retrieved after prior context testing.

Altogether, the hyper-reactivation of the ACC engram seems to be specific to the PTSD-like memory profile whatever the order of testing.

### PTSD-like memory impairments persist over remote time and remain associated with ACC engram reactivation

PTSD is a long-lasting disorder, raising the question of whether the memory impairments induced by corticosterone in our model persist over remote time. To address this, TRAP2;Ai14 mice were conditioned as described above, injected with either corticosterone or saline together with 4-OHT after conditioning, and then tested 30 days later in the tone and context retrieval sessions (Fig. 2a). After this 30-day delay, Cort-injected mice still displayed an enhanced response to the trauma-associated tone, as shown by a significantly increased tone ratio compared with Sal controls (Fig. 2b,c). Interestingly, Sal-injected mice showed elevated freezing before tone presentation in the neutral context, suggesting some degree of fear generalization at this remote time point. Despite this, Cort-injected mice retained a selective hyper-responsiveness to the tone. When re-exposed to the conditioning context the following day, Cort-injected mice showed impaired contextual memory, with significantly reduced freezing during the first two minutes of the test compared with Sal controls (Fig. 2d,e). Consistently, the integrated PTSD-like score was significantly higher in Cort-injected mice compared to controls (Fig. 2f). These results indicate that PTSD-like memory is not transient, but persists over remote time.

**Fig. 2.**
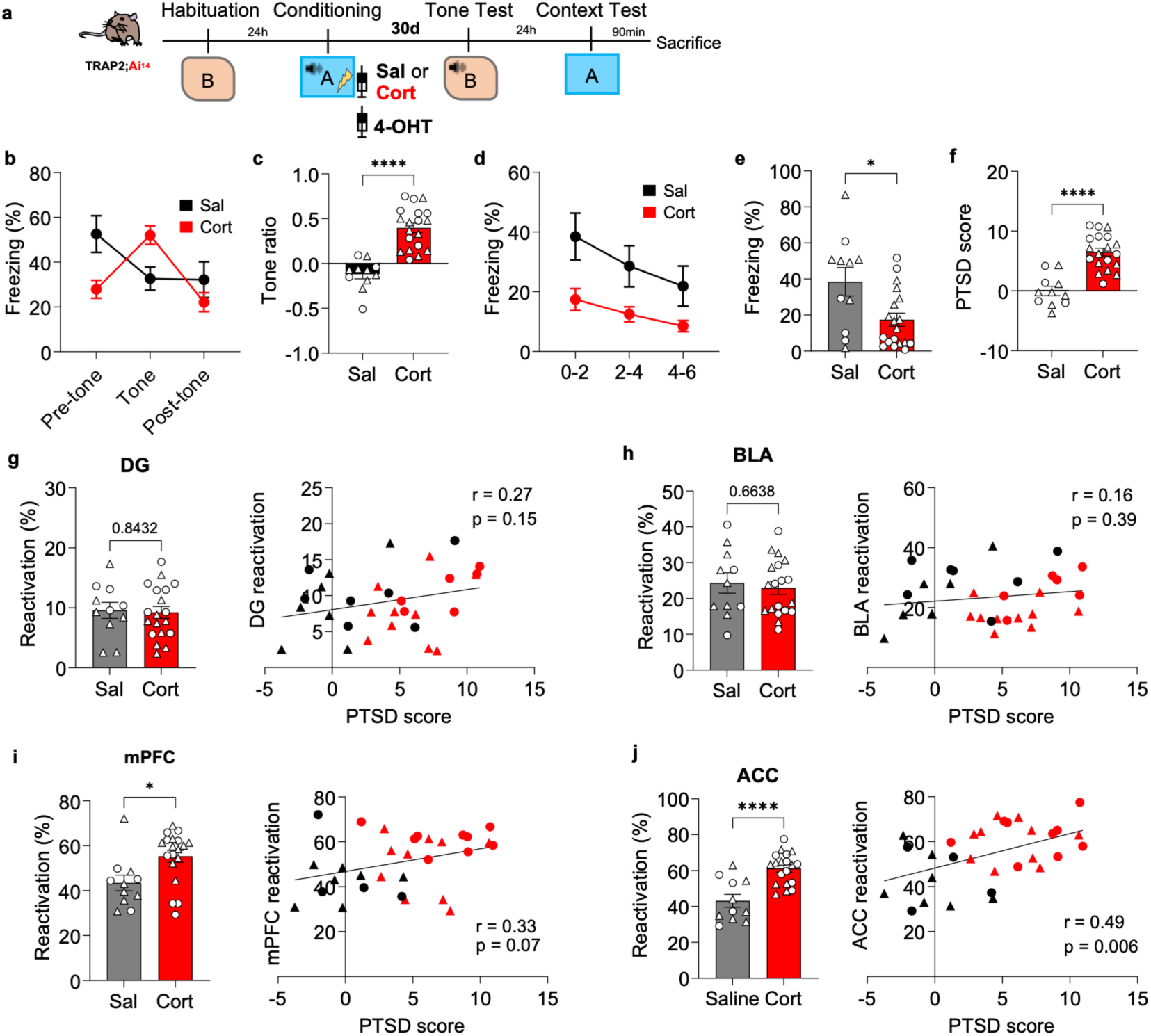
Traumatic memory profiles are preserved across remote timepoints, characterized by elevated and behaviorally correlated ACC engram reactivation. **a**, Experimental protocol for remote memory assessment. *Trap2;Ai14* mice were habituated to a neutral context (CtxB). The following day, mice underwent fear conditioning in CtxA, immediately followed by either Corticosterone (Cort, *n =* 19) or Saline (Sal, *n* = 11) injections combined with 4-OHT administration. Mice were tested 30 days later in CtxB in the presence of the tone, and evaluated the following day in the original conditioning context (CtxA). **b**, Percentage of freezing in CtxB. Cort mice exhibited an elevated fear response during tone presentation compared to Sal mice *(p* < 0.05). **c**, Tone ratio, which was significantly higher in Cort mice *(****p* < 0.0001). **d**, Percentage of freezing in CtxA. Cort mice displayed a decreased total freezing level compared to Sal mice *(p* < 0.01, main effect of group). **e**, Freezing response during the first two minutes in CtxA, which was significantly lower in Cort mice (p < 0.05). **f**, PTSD score, showing a marked increase in Cort mice compared to Sal mice (****p < 0.0001). **g, h**, Engram reactivation in the dentate gyrus (DG) (g) and basolateral amygdala (BLA) (h). Reactivation rates did not differ between Cort and Sal groups, and did not correlate with PTSD scores. **i**, Engram reactivation in the medial prefrontal cortex (mPFC). Reactivation was significantly higher in Cort mice, showing a strong trend toward correlating with individual PTSD scores. **j**, Engram reactivation in the anterior cingulate cortex (ACC). Reactivation was significantly elevated in Cort mice and positively correlated with PTSD scores. (For statistical details see table 1).

We then assessed whether remote PTSD-like memory was associated with persistent alterations in engram reactivation across the fear-memory network. At this time point, DG and BLA engram reactivation no longer differed between Cort-and Sal-injected mice and did not significantly correlate with the PTSD-like score (Fig. 2g,h). In contrast, both mPFC and ACC showed increased engram reactivation in Cort-injected mice (Fig. 2i,j). However, only ACC reactivation was significantly correlated with the PTSD-like score, whereas mPFC reactivation showed only a positive trend. Together, these findings suggest that, over remote time, PTSD-like memory becomes particularly associated with persistent ACC engram reactivation, supporting a role for the ACC in the long-term maintenance of maladaptive traumatic memory.

### Inhibition of ACC engram cells prevents PTSD-like memory expression and normalizes downstream engram reactivation

Given the enhanced reactivation of ACC engram cells and its positive correlation with the PTSD-like score, we next asked whether this cortical engram population was functionally required for the expression of PTSD-like memory impairments. To selectively inhibit ACC neurons recruited during traumatic memory encoding, AAV-DIO-mCherry-hM4Di or a control AAV-DIO-mCherry virus were bilaterally injected into the ACC of TRAP2 mice. After viral expression, mice underwent the conditioning protocol and received 4-OHT immediately after conditioning, allowing Cre-dependent expression of the inhibitory DREADD hM4Di specifically in conditioning-activated ACC neurons, or its control virus. The tagged ACC engram cells were then transiently inhibited by systemic injection of the DREADD agonist DCZ 45 min before each retrieval session, first before tone testing and then before context testing the following day (Fig. 3a,b). As expected, Cort-injected mice expressing the control virus showed increased freezing during tone presentation, confirming hyper-responsiveness to the trauma-related but irrelevant cue. In contrast, Cort-injected mice in which ACC engram cells were inhibited no longer displayed a marked freezing response to the tone (Fig. 3c). Indeed, the tone ratio remained elevated in control-virus Cort mice but was normalized in hM4Di Cort mice to levels comparable to Sal controls (Fig. 3d). Thus, acute inhibition of ACC engram cells prevented the expression of trauma-related cue hypermnesia.

**Fig. 3.**
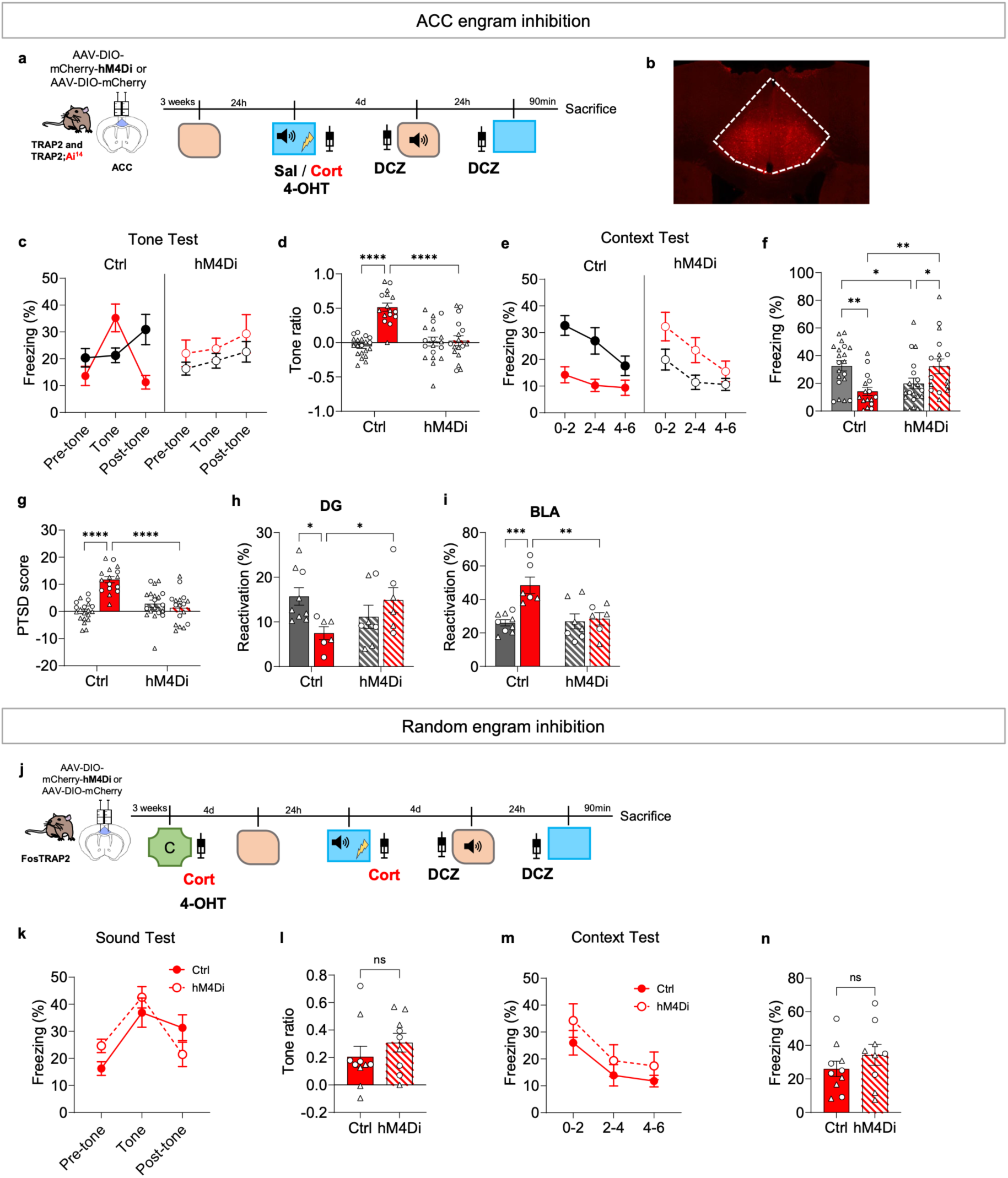
Chemogenetic inhibition of ACC engram cells—but not a randomly activated ACC population—restores the traumatic PTSD-like memory profile to an adaptive state. ACC Engram Inhibition **a**, Experimental protocol for fear-engram inhibition. *Trap2;Ai14* or *Trap2* mice were microinjected with either an inhibitory virus (AAV-DIO-hM4Di-mCherry) or a control virus (AAV-DIO-mCherry) into the ACC. Three weeks later, mice were habituated to a neutral context (CtxB). The following day, they underwent fear conditioning in CtxA, immediately followed by Corticosterone (Cort) or Saline (Sal) injections and 4-OHT administration. Mice were tested 4 days later in a neutral context (CtxB) with tone presentation, and evaluated the following day in CtxA. Deschloroclozapine (DCZ) was administered 45 min prior to each behavioral test. **b**, Representative histological illustration of virus infusion targeting the ACC. **c, d**, Freezing behavior in response to the tone in CtxB. In control-virus mice (Ctrl), Cort mice displayed elevated freezing compared to Sal mice, shown across the 6­min session (c) and quantified via tone ratio (d). Conversely, inhibition of the ACC engram *(hM4Di)* prevented this elevated fear response and high tone ratio in Cort mice. **e, f**, Freezing behavior in the conditioning context (CtxA). In Ctrl mice, Cort animals showed reduced overall freezing across the 6-min session (e) and significantly lower freezing during the first two minutes (f) compared to Sal controls. ACC engram inhibition normalized this deficit, restoring Cort-hM4Di freezing to control levels. **g**, Calculated PTSD scores, demonstrating that the elevated score in Cort-Ctrl mice is significantly rescued by ACC engram inhibition. **h, i**, Engram reactivation rates in the DG (h) and BLA (i). ACC engram inhibition normalized downstream engram reactivation deficits in Cort mice. Data are represented as mean ± SEM; *\*p* < 0.05, *\*\*p* < 0.01, *\*\*\*p* < 0.001, *\*\*\*\*p* < 0.0001. *Cort: n = 34 total (Ctrl: n = 6 Trap2;AH4, n = 11 Trap2; hM4Di: n = 6 Trap2;AH4, n = 11 Trap2). Sal: n = 39 total (Ctrl: n = 9 Trap2;AH4, n = 11 Trap2; hM4Di: n = 7 Trap2;AH4, n = 12 Trap2).* (For statistical details see table 1). Random ACC Population Inhibition Control **j**, Experimental protocol for the inhibition of a non-specific, randomly activated ACC cell population. *FosTrap2* mice were injected with AAV-DIO-hM4Di-mCherry *(n =* 9) or AAV-DIO-mCherry control *(n =* 10) in the ACC. Mice were exposed to Context C paired with Cort and 4-OHT to label a random cell population. Four days later, all mice were fear-conditioned in CtxA and given a Cort injection. Testing in CtxB (with tone) and CtxA occurred 4 days later, with DCZ administered 45 min prior to testing, k-n, Behavioral outcomes following random ACC cell inhibition. Ctrl and hM4Di groups showed no significant differences in tone-induced freezing in CtxB (k), tone ratio (I), total context-induced freezing in CtxA (m), or context-induced freezing during the first two minutes (n).

We then examined contextual memory recall. In control virus-injected mice, Cort treatment induced the expected contextual memory impairment, with reduced freezing upon re-exposure to the conditioning context. Inhibiting ACC engram cells before the context test prevented this contextual memory deficit as hM4Di Cort mice showed restored freezing levels compared with control Cort mice (Fig. 3e,f). Interestingly, ACC engram inhibition also reduced contextual freezing in Sal mice, consistent with a contribution of ACC engram cells to normal contextual fear memory expression. When both behavioural dimensions were integrated into the PTSD-like score, hM4Di Cort mice no longer showed the elevated score observed in control Cort mice, indicating a normalization of the PTSD-like memory profile (Fig. 3g).

Finally, we asked whether inhibiting the ACC engram could also normalize the altered reactivation of other memory-related regions identified in Figure 1. To address this, the same strategy was applied in TRAP2;Ai14 mice, allowing quantification of tdTomato-tagged engram reactivation in the DG and BLA after retrieval. In control Cort mice, DG reactivation was reduced and BLA reactivation was increased, reproducing the abnormal engram reactivation pattern observed previously. In contrast, inhibition of ACC engram cells normalized tdTomato-cell reactivation in both DG and BLA (Fig. 3h,i). These results suggest that ACC engram activity is not only required for the behavioural expression of PTSD-like memory, but also contributes to the network-level imbalance of engram reactivation across hippocampal and amygdalar structures.

### The effect of ACC engram inhibition is specific to neurons recruited during traumatic memory encoding

To clarify the nature of this inhibitory effect, we addressed whether it was truly engram-specific. Because corticosterone can enhance neuronal activation (Jaszczyk and Juszczak, 2021), and previous studies have shown that neurons with higher activity become allocated to the engram (Zhou et al., 2009), the pronounced behavioral effect observed after ACC engram inhibition might have stemmed from a biased and enhanced iDREADD expression in these Cort activated ACC neurons, rather than the targeted suppression of neurons engaged during traumatic encoding. To address this, we tested whether inhibiting a Cort-and novelty-activated ACC neuronal population recruited outside the conditioning episode would similarly impair PTSD-like memory expression. TRAP2 mice were injected with either AAV-DIO-mCherry-hM4Di or the control virus into the ACC. Following viral expression, mice were exposed to a neutral context (context C) for 3 min and treated with corticosterone together with 4-OHT, thereby tagging a Cort-activated ACC neuronal population unrelated to trauma. Four days later, mice underwent the standard traumatic conditioning protocol, including post-conditioning Cort. The previously tagged neutral-context ACC population was then inhibited with DCZ prior to tone and context retrieval sessions (Fig. 3j,k).

In contrast to the effect of silencing conditioning-recruited cells, inhibiting this neutral-context, Cort-activated ACC population failed to prevent PTSD-like memory expression. Cort-injected mice still exhibited heightened freezing during tone presentation, with no significant difference in tone ratio between hM4Di and control groups (Fig. 3l,m). Similarly, inhibiting this non-conditioning ACC population did not alter freezing behaviour during re-exposure to the conditioning context (Fig. 3n,o). Together, these results demonstrate that the behavioural rescue observed after ACC engram inhibition is not a non-specific by-product of suppressing any Cort-or novelty-activated population, but rather depends strictly on targeting the neurons recruited during traumatic memory encoding.

To further control for the specificity of this effect, we also tagged an ACC neuronal population activated by corticosterone in the home cage before conditioning. As in the neutral-context experiment, inhibition of this home-cage Cort-activated population during subsequent retrieval did not modify freezing responses to the trauma-associated tone or to the conditioning context (Suppl Fig. 3a–e). Together, these control experiments demonstrate that the ACC population driving PTSD-like memory expression is specifically recruited during traumatic memory encoding, rather than reflecting a nonspecific Cort-responsive neuronal ensemble.

## DISCUSSION

In this study, we identify a cortical engram mechanism supporting PTSD-like memory impairments in a mouse model combining trauma-related cue hypermnesia and contextual amnesia. This behavioural dissociation is highly relevant to PTSD, which is defined not merely by heightened fear, but by intrusive trauma-related responses coupled with an inability to contextualize the traumatic event (Brewin et al., 1996; Liberzon and Abelson, 2016; Al Abed et al., 2020; Alexandra Kredlow et al., 2022). Notably, this behavioural dissociation was observed for the first time in female mice. Incorporating both sexes is crucial in preclinical trauma research, as women are two to three time more likely than men to develop PTSD following a traumatic event (Olff, 2017). Using an activity-dependent tagging strategy based on the TRAP approach, which allows genetic access to neurons activated during a defined temporal window (Guenthner et al., 2013; DeNardo et al., 2019), we show that corticosterone administered after fear conditioning does not simply amplify the overall neuronal recruitment. Rather, it reshapes the pattern of engram reactivation across a distributed fear-memory network. At recent time points, PTSD-like memory was associated with reduced reactivation of DG engram cells and enhanced reactivation of BLA and ACC engram cells. Importantly, BLA and ACC reactivation positively correlated with the severity of the PTSD-like behavioural phenotype, suggesting that maladaptive memory expression emerges from an imbalance between hippocampal and cortico-amygdalar components of the memory trace.

This interpretation fits with the broader engram framework, in which memories are supported by specific neuronal populations recruited during learning and reactivated during recall (Liu et al., 2012; Josselyn and Tonegawa, 2020). However, our results extend this framework to a PTSD-like state by showing that maladaptive memory is not explained by a uniform increase in engram size or global neuronal activation. Instead, traumatic memory appears to involve an abnormal redistribution of engram reactivation across regions involved in contextual discrimination, emotional salience, and cortical control. The reduced DG reactivation observed in corticosterone-treated mice may reflect impaired access to a precise contextual representation, consistent with the role of the hippocampus and dentate gyrus in contextual processing and fear generalization (Liberzon and Abelson, 2016; Bian et al., 2019). Conversely, increased BLA reactivation is consistent with the central role of the amygdala in assigning emotional significance to aversive cues and supporting fear-memory expression (Mahan and Ressler, 2012; Sun et al., 2020), and extends the previous literature on the increased engram in lateral amygdala of a Cort-induced generalized fear memory (Lesuis et al., 2025).

A major finding of this work is that the ACC engram is not only correlated with PTSD-like memory impairments, but is functionally required for their expression. Chemogenetic inhibition of ACC neurons recruited during traumatic memory encoding abolished hyper-responsiveness to the trauma-associated tone and restored contextual freezing in corticosterone-treated mice. This dual rescue is particularly important because the PTSD-like phenotype in our model is defined by the coexistence of two apparently opposite symptoms: excessive response to a trauma-related but non-predictive cue, and impaired memory for the predictive conditioning context. The ACC has been repeatedly implicated in remote contextual fear memory, threat processing, and generalized fear responses (Frankland, 2004; Bian et al., 2019; Alexandra Kredlow et al., 2022). Our findings add a causal engram-level mechanism by showing that ACC neurons recruited during traumatic encoding can drive both cue-related hypermnesia and contextual memory impairment.

Our data further indicate that ACC engram activity contributes to the broader network imbalance observed during PTSD-like memory recall. Inhibition of conditioning-recruited ACC engram cells normalized the abnormal reactivation of both DG and BLA engrams. This suggests that ACC activity may exert top-down control over hippocampal and amygdalar memory components, biasing recall toward trauma-associated cue reactivity while weakening contextual memory retrieval. This interpretation is consistent with circuit studies showing that ACC projections can influence fear generalization and amygdala-dependent defensive responses (Jhang et al., 2018; Bian et al., 2019; Ortiz et al., 2019). In this framework, the ACC may act as a pathological hub coordinating two core dimensions of PTSD-like memory: excessive emotional responding and impaired contextualization.

The specificity of this mechanism is supported by the control experiments in which inhibition of ACC neurons activated by corticosterone outside the conditioning episode failed to modify PTSD-like memory expression. Inhibiting a corticosterone-activated ACC population tagged either in a neutral context or in the home cage did not prevent tone hyper-responsiveness or restore contextual freezing. These findings argue against a nonspecific effect of suppressing any corticosterone-responsive ACC population. Instead, they indicate that the behavioural rescue depends on inhibiting ACC neurons specifically recruited during traumatic memory encoding. This distinction is essential, because glucocorticoids can broadly modulate neuronal excitability, stress-related memory encoding, and activity-dependent neuronal recruitment. Thus, the ACC population driving PTSD-like memory expression appears to be defined by the conjunction of traumatic encoding and corticosterone exposure, rather than by corticosterone responsiveness alone.

An additional important observation is that the PTSD-like phenotype persists over remote time. Thirty days after conditioning, corticosterone-treated mice still displayed enhanced responding to the trauma-associated tone, reduced contextual freezing, and an elevated PTSD-like score. At this remote time point, DG and BLA engram reactivation no longer differed between groups (Cort-vs. Sal-injected mice), whereas ACC reactivation remained increased and continued to correlate with the severity of PTSD-like behaviour. This temporal shift is consistent with systems consolidation models in which cortical networks, including the ACC and prefrontal cortex, become progressively more engaged in remote memory storage and retrieval (Frankland, 2004; Kitamura et al., 2017; DeNardo et al., 2019; Lee et al., 2023). Our results extend this concept by showing that compared to adaptive fear memory, a maladaptive traumatic memory state can remain anchored to persistent and stronger ACC engram reactivation.

These findings also have implications for how PTSD-like memory should be conceptualized experimentally. Many preclinical studies equate strong fear memory with traumatic or PTSD-like memory. However, PTSD involves a pathological organization of memory, including intrusive responses to trauma-related cues and impaired contextualization, rather than simply stronger fear expression. Our model captures this dissociation by combining cue hyper-responsiveness with contextual memory impairment. The present data suggest that this behavioural dissociation is mirrored at the engram level by an abnormal redistribution of memory reactivation, rather than by a uniform expansion of the fear engram. This point is important because it indicates that maladaptive memory may arise not from the amount of neuronal recruitment during encoding, but from the way specific memory ensembles are re-engaged and coordinated during recall.

Several limitations of the present study should be considered. First, while cFos-based tagging and recall mapping provide powerful access to activity-dependent neuronal ensembles, they do not capture the full dimensions of neuronal coding, synaptic plasticity, or real-time firing dynamics. Future studies utilizing calcium imaging or electrophysiological approaches will be required to determine how ACC engram cells precisely encode trauma-related cue information, contextual representations, and their interaction during recall. Second, although chemogenetic inhibition demonstrates a causal role for ACC engram cells in memory expression, it does not fully elucidate the downstream projection targets, specific neuronal subtypes, or local circuit mechanisms involved. Given the established role of ACC–amygdala interactions in driving fear and defensive responses as well as generalization (Jhang et al., 2018; Ortiz et al., 2019), the ACC-to-BLA pathway represents a strong candidate circuit; however, additional projections to hippocampal, thalamic, or other prefrontal regions may also contribute to these phenotypes. Crucially, inputs from the ACC to BLA are predominantly glutamatergic/excitatory, yet they typically yield net BLA inhibition due to the recruitment of local feedforward inhibiting microcircuits (McGarry and Carter, 2016; Jhang et al., 2018). In our model, however, ACC hyperactivation appears to coexist with BLA hyperactivation. This paradox suggests that the canonical ACC-to-BLA feed-forward inhibition pathway may be dysfunctional in this context, ultimately driving maladaptive network recruitment and the breakdown of emotional regulation.

Finally, while the animal model used is the first to capture the cardinal feature of PTSD, namely the paradoxical alteration of memory, and is extremely relevant for identifying a specific memory engram of PTSD (vs. adaptive fear memory), it does not recapitulate all symptoms of PTSD. Using alternative models in the future could help complement this study.

In conclusion, our study identifies ACC engram cells as a causal substrate for PTSD-like memory impairments. Rather than supporting the view that traumatic memory is merely a stronger fear memory, our findings suggest that PTSD-like memory emerges from a pathological reorganization of engram reactivation across hippocampal, amygdalar, and cortical regions. The ACC appears to occupy a central position in this network, driving both trauma-related cue hypermnesia and contextual memory impairment, while also contributing to the long-term persistence of the maladaptive memory state. Targeting the mechanisms that stabilize or reactivate ACC traumatic-memory ensembles may therefore represent a promising strategy to restore adaptive memory processing after trauma.

## METHODS

### Mice

Male and female FosTRAP (TRAP2) mice (B6.129(Cg)-^Fostm1.1(cre/ERT2)Luo^/J) and FosTrap;Ai^14^ (B6.129(Cg)-^Fostm1.1(cre/ERT2)Luo^/J X B6.Cg-Gt(ROSA)26Sor^tm9(CAG-tdTomato)Hze^/J) (TRAP2,Ai^14)^ were used for this work. They were housed in groups of 4 to 5 animals or by pairs during the behavioral experiments with food and water ad libitum and were maintained in a room under controlled light and dark cycle (12/12 h; light starts at 7:00 a.m.), temperature (22 ± 2 C), and humidity (55 ± 10%). Adult mice (aged between 10 weeks and 15 weeks) were used. All experiments were performed in accordance with the European Directive for the care and use of laboratory animals (2010-63-EU) and the animals care guidelines issued by the animal experimental committee of Bordeaux University (CE50, agreement number A33-063-100; authorizations No. 56762; No. 38389).

### Fear conditioning procedure

The day before fear conditioning, all mice were individually placed for 2,5 min into a chamber (30 × 30 × 40 cm) with an opaque PVC floor, with a brightness of 20 lux. The box was cleaned with 1% acetic acid before each trial. This pre-exposure allowed the mice to acclimate and become familiar with the chamber later used for the tone re-exposure test. Acquisition of fear conditioning was performed in the conditioning context, (30 × 30 × 40cm), in a brightness of 80 lux, given access to the different visual-spatial cues located on the walls of the box. The floor of the chamber consisted of 20 stainless-steel rods (6mm diameter), spaced 1 cm apart and connected to a shock generator. The box was cleaned with 70% ethanol before each trial. All animals were trained with a tone–shock unpairing procedure, meaning that the tone was non-predictive of the foot-shock. This training procedure, routinely used in our laboratory, promotes the processing of contextual cues in the foreground. Briefly, each animal was placed in the conditioning chamber for approximately 4min during which it received two tone cues (65 dB, 1 kHz, 15 s) and two foot-shocks (squared signal: 0.4mA, 50Hz, 1 s) according to the following distribution: 100 s after being placed in the chamber, animals received a shock, then, after a 30 s interval, a tone; finally, after a 20 s delay, the same tone and the same shock spaced by a 30 s interval were presented. After 20 s, animals were returned to their home cage. In this tone–shock unpairing procedure, as the tone is never immediately followed by shock delivery, animals identify the conditioning context (set of static background contextual cues that constitutes the environment in which the conditioning takes place), and not the tone, as the right predictor of the shock (Kaouane et al., 2012).

Four days and five days after the acquisition of fear conditioning, mice were submitted to 2 memory retention tests, to the tone presentation and to the context presentation respectively. During these two memory tests, animals were continuously recorded for off-line manual scoring of freezing by an observer blind of experimental groups. Freezing behavior of animals, defined as a lack of all movement except for respiratory-related movements, was used as an index of conditioned fear response. Mice were first submitted to the tone re-exposure test in the safe familiar chamber during which three successive recording sessions of the behavioral responses were performed: one before (first 2min, “pre-tone”), one during (next 2 min, “tone”), and one after (2 last min, “post-tone”) tone presentation. Conditioned response to the tone is expressed by the percentage of freezing during the tone presentation compared to the levels of freezing expressed before and after tone presentation. The strength and specificity of this conditioned fear is attested by a ratio that considers the percentage of freezing increase to the tone with respect to a baseline freezing level (i.e., pre-and post-tone periods mean). The tone ratio is calculated as follows: [% freezing during tone presentation−(% pre-tone period freezing + % post-tone period freezing)/ 2] / [% freezing during tone presentation + (% pre-tone period freezing +% post-tone period freezing)/ 2]. The following day, mice were submitted to the context re-exposure test: they were placed for 6min in the conditioning chamber. Freezing to the context was calculated as the percentage of the total time spent freezing during the successive three blocks of 2-min periods of the test.

### PTSD score calculation

A Z-score representing the intensity of PTSD-like memory symptoms, and integrating both the contextual and the cued responses, was calculated for each mouse. Z scores values were calculated individually for the tone test and the context test using the following parameters: for tone test, X = ratio; for context test, X = Freezing max 0-2 min – Freezing 0-2 min. Z score values = (X-μ)/ σ where μ is the mean and σ the standard deviation for the control group. PTSD score was calculated as (Z tone + Z context)/ 2.

### 4-Hydroxytamoxifen (4-OHT) preparation and systemic injection

4-hydroxytamoxifen (4-OHT; Hello Bio, Cat# HB6040) was dissolved at 20 mg/mL in ethanol by shaking at 37°C for 15 min and was then aliquoted and stored at –20°C for up to several weeks. Before use, 4-OHT was mixed with a 1:4 mixture of castor oil: sunflower seed oil (Sigma, Cat # S259853 and S5007) to give a final concentration of 10 mg/mL 4-OHT, and the ethanol was evaporated by shaking at 65°C for 1 hour. The final 4-OHT solutions were always used on the day they were prepared. All injections were delivered intraperitoneally (i.p.) after fear conditioning to achieve a final concentration of 50 mg/kg of 4-OHT to mice. After 4-OHT administration, animals were returned to their home cages.

### Systemic injection of corticosterone

Corticosterone (2-hydroxypropyl-β-cyclodextrin complex; 2.5 mg/kg in a volume of 0.1 ml/10 g bodyweight) or vehicle (NaCl 0.9%) was administered i.p. immediately after the acquisition of fear conditioning. After the injection, animals were returned to their home cage. The dose of corticosterone was selected on the basis of previous results indicating that such dose (i) is in the range of concentrations induced by stress in the plasma and (ii) effectively induces PTSD-like memory in mice when combined with fear conditioning.

### Systemic injection of DCZ

DCZ (Deschloroclozapine dihydrochloride, Hello Bio, #HB9126) was dissolved in NaCl and injected at the concentration of 0,1mg/Kg.

### Surgery

Mice were anaesthetized with 4% isoflurane for 5min and placed in the stereotaxic frame, where they were maintained on 1-2% isoflurane for the duration of the surgery. Analgesia was provided by a subcutaneous (SC) injection of Metacam (1 mg/kg) and hydration ensured by NaCl 0.9% SC injection. All viral injections were performed using pulled glass pipettes (WPI) and a microinjector (Nanoliter2020), set to a delivery rate of 50nL/min, into the anterior cingulate cortex (AP +0.6, ML +-0.3, DV-1.8 from bregma). Pipettes were left in place for 5 additional minutes to prevent virus spreading. Mice were given metacam and monitored for the 3 consecutive days after surgery.

### Adeno-associated virus (AAV)

The viruses used were purchased premade (ZTH #v84-8, ssAAV-8/2-hSyn1-dlox-hM4D(Gi)_mCherry(rev)-dlox-WPRE-hGHp(A), ZTH #116-8 ssAAV-8/2-hSyn1-dlox-mCherry(rev)-dlox-WPRE-hGHp(A)). Used in TRAP2 or TRAP2;Ai14 mice, these virus allowed for the expression of an inhibitory DREADD (designer receptors activated by designer drugs) hM4Di or its’ control (same construct without hM4Di) in activated cells.

### Brain slicing and immunohistochemistry

Animals were perfused transcardially with a phosphate-buffered solution of 4% paraformaldehyde. After at least 3 days of fixation, brains were cut with a vibratome at 40-μm thickness. 1-in-6 free-floating sections were processed according to a standard immunohistochemical procedure. Briefly, for fluorescence labeling, slices were washed with PBS (0.1M) four times for 10 before being incubated in blocking solution (5% normal goat serum or 5% normal donkey serum in 0,3% Triton X-100-PBS). They were then incubated with primary antibody rabbit anti c-Fos antibody (1:5000, Synaptic Systems, #226008) or goat anti RFP antibody (1:1000, Rockland, #039200-101-379) for 48h at 4°C under agitation followed by secondary antibody incubation (Goat anti-rabbit Alexa488, 1:1000, Invitrogen, #A-11008, donkey anti goat A568, 1:1000 #A-11057) for 2h at room temperature, also under agitation. DAPI (Sigma Aldrich, #D9542) allowed for cell nuclei detection.

### Microscopy data analysis

Slices were mounted on glass coverslips and imaged in a slide scanner Hamamatsu Nanozoomer 2.0HT with a 20x objective for cFos and tdTomato quantification or in a confocal Leica DM6000 TCS SP5 MP for ACC reactivation and Leica DM5500 TCS SPE for DG and BLA reactivation. For ACC reactivation, 2 slices per mouse were imaged, acquiring a Z-stack of 4um at 1024×1024 resolution, with a 20x oil objective. Microscopy images were analysed using Qupath.

## Statistical Analysis

All statistical analysis were used GraphPad Prism 11. For comparisons between two groups, unpaired t-tests were used as appropriate. For comparisons among more than two groups, one way, two-way or three-way repeated measures ANOVA was performed. When significant, ANOVA was followed by posthoc analyses.

