## Supplementary figures for "Anterior cingulate cortex engram drives Post traumatic stress disorder-like memory impairments in a rodent model of traumatic fear"

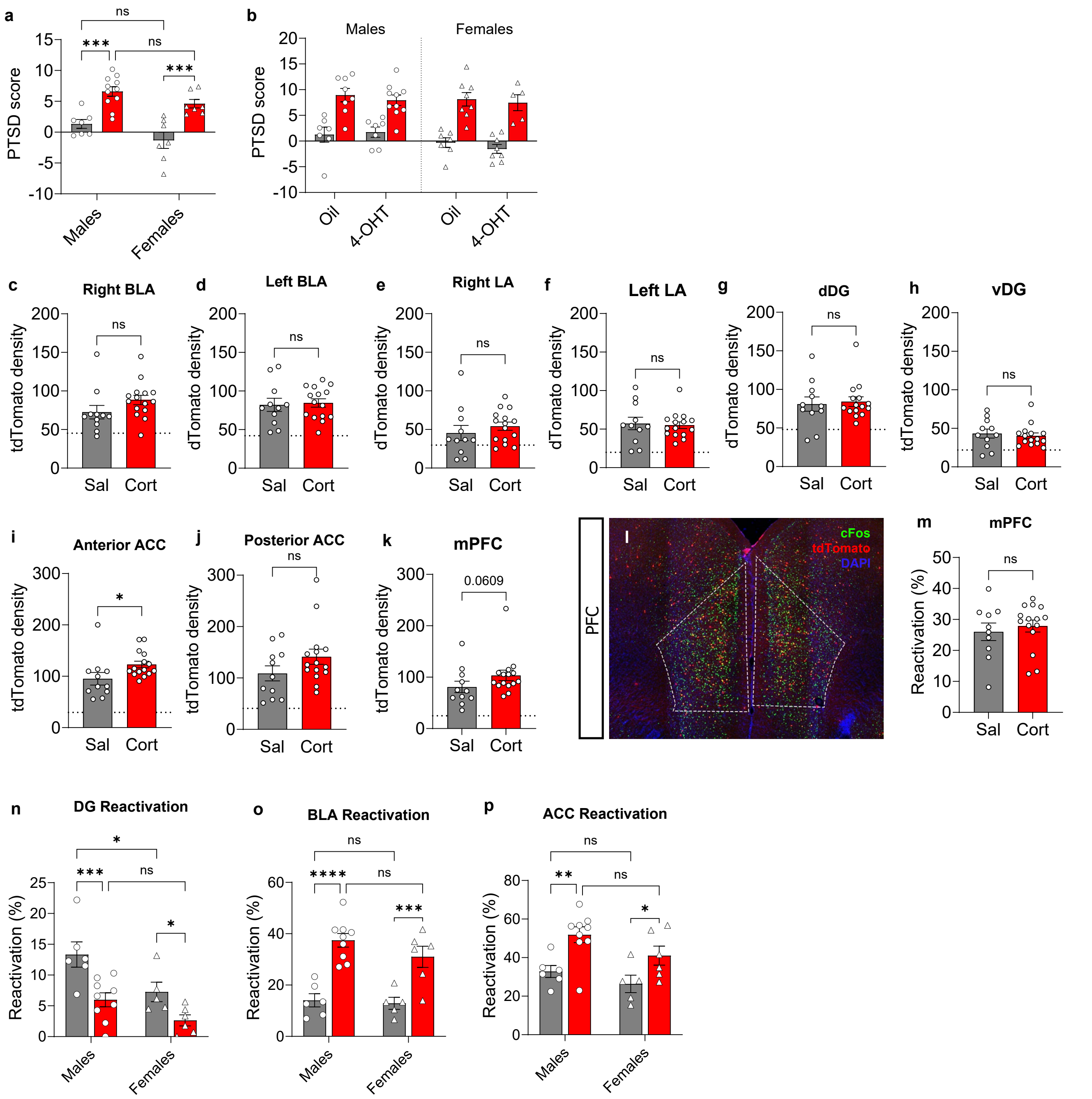


**Suppl Fig 1**

a. No differences were found between males and females : for tone ratio in Ctx B, percentage of freezing in Ctx A in Cort mice – only Sal females froze more in Cxt A- and for PTSD score.

b. No difference was found in PTSD score between male and female mice injected with 4-OHT and oil (vehicle). (****$p < 0.0001$; ***$p < 0.001$; **$p < 0.01$; *$p < 0.05$)

**c-k** Quantified tdTomato+ engram cell density: Densities did not significantly differ between groups in the right BLA, left BLA, Right LA, Left LA, dorsal DG, ventral DG, posterior ACC and mPFC. tdTomato density was higher in Cort mice in the anterior ACC (dashed lines represent baseline activation) *$p < 0.05$.

**l,m** Representative image of mPFC. No differences were found in mPFC reactivation.

**n-p** DG tdTomato+ cells showed significantly decreased reactivation in Cort mice during recall in both males and females. Cort mice displayed significantly higher reactivation of tdTomato+ cells compared to Sal mice in BLA and ACC both in males and females.

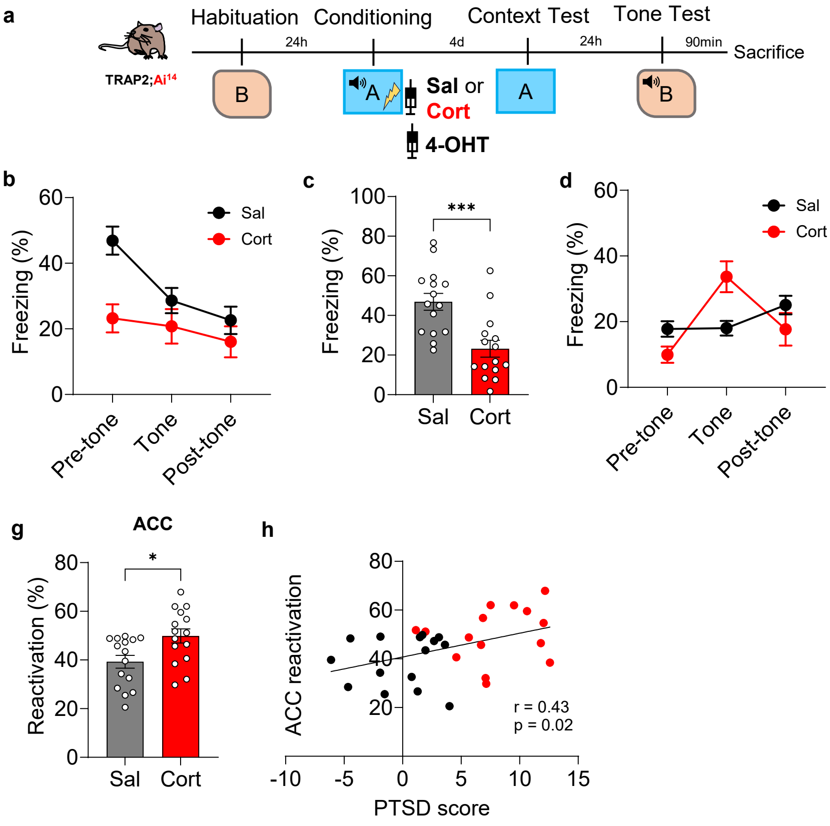


**Suppl Fig. 2. Traumatic fear is expressed after inverting the order of the retrieval session and induces hyper-reactivation of engram cells in the ACC.**

**a,** Experimental protocol for recent memory assessment. *Trap2;Ai14* mice were habituated to a neutral context (CtxB). The following day, mice were conditioned in CtxA and immediately received either Corticosterone (Cort, $n$ = 15) or Saline (Sal, $n$ = 15) injections combined with 4-OHT administration. Mice were tested 4 days later in CtxA, and evaluated the following day in the CtxB in the presence of the tone.

**b,** Percentage of freezing in CtxA. Cort mice displayed a decreased total freezing level compared to Sal mice (***$p$ < 0.001, main effect of group).

**c,** Freezing response during the first two minutes in CtxA, which was significantly lower in Cort mice (***$p$ < 0.001).

**d,** Percentage of freezing in CtxB. Cort mice exhibited an elevated fear response during tone presentation compared to Sal mice (***$p$ < 0.001).

**e,** Tone ratio, which was significantly higher in Cort mice (****$p$ < 0.0001).

**f,** Calculated PTSD scores, showing a marked increase in Cort mice compared to Sal mice (****$p$ < 0.0001).

**g, h** Engram reactivation and behavioral correlations in ACC. Cort mice displayed significantly higher reactivation of tdTomato+ cells compared to Sal mice, which positively correlated with individual PTSD scores (*$p$ < 0.05).

others

(For statistical details see table 1).

HC random population inhibition

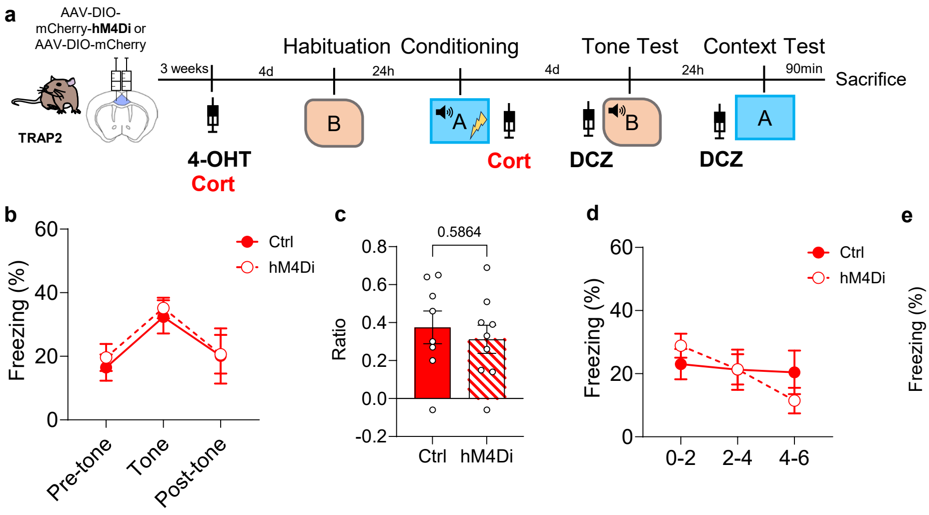


**Suppl Fig. 3. Inhibiting a home-cage Cort-activated population does not alter traumatic memory.**
**a,** Experimental protocol for the inhibition of a home-cage (HC) non-specific ACC population. *FosTrap2* mice were injected with AAV-DIO-hM4Di-mCherry (*n* = 9) or AAV-DIO-mCherry control (*n* = 8) in the ACC. Mice were injected with Cort and 4-OHT in their HC to label a random cell population. Four days later, all mice were habituated as usual in CtxB and one day after fear-conditioned in CtxA and given another Cort injection. Testing in CtxB (with tone) and CtxA occurred 4 days later, with DCZ administered 45 min prior to testing. **b-e,** Behavioral outcomes following random ACC cell inhibition. Ctrl and hM4Di groups showed no significant differences in tone-induced freezing in CtxB (**b**), tone ratio (**c**), total context-induced freezing in CtxA (**d**), or context-induced freezing during the first two minutes (**e**).

| Figure | Panel | Statistical test | Effect | Statistic | p-value |
|---|---|---|---|---|---|
| 1 | c | Two-way ANOVA (repeated measures) | Group | F (1,30)= 2.47 | 0.13 |
| | | | Time | F (2,60)= 3.83 | 0.03 |
| | | | Group x Time | F (2,60) = 18.57 | <0.0001 |
| | d | Unpaired t-test | | | <0.0001 |
| | e | Two-way ANOVA (repeated measures) | Group | F (1,30)= 15.71 | 0.0004 |
| | | | Time | F(2,60) = 14.69 | <0.0001 |
| | | | Group x Time | F (2,60) = 0.32 | <0.0001 |
| | f | Unpaired t-test | | t (30) = 3.70 | 0.0009 |
| | g | Unpaired t-test | | t (30) = 6.04 | <0.0001 |
| | i | Mann-Whitney | | U = 69 | 0.51 |
| | j | Unpaired t-test | | t (24) = 0.59 | 0.56 |
| | k | Unpaired t-test | | t (24) = 3.51 | 0.0018 |
| | l | Linear regression | | r= -0.27 | 0.18 |
| | n | Unpaired t-test | | t (24) = 1.01 | 0.32 |
| | o | Mann-Whitney | | U = 72 | 0.61 |
| | p | Unpaired t-test | | t (24) = 6.855 | <0.0001 |
| | q | Linear regression | | r=0.71 | <0.0001 |
| | s | Mann-Whitney | | U = 47 | 0.069 |
| | t | Unpaired t-test | | t (24) = 0.73 | 0.47 |
| | u | Unpaired t-test | | t (24) = 3.86 | 0.0008 |
| | v | Linear regression | | r=0.43 | 0.03 |

| Figure | Panel | Statistical test | Effect | Statistic | p-value |
|---|---|---|---|---|---|
| 2 | b | Two-way ANOVA (repeated measures) | Group | F (1,28) = 0.62 | 0.43 |
| | | | Time | F (2,56) = 10.22 | 0.0002 |
| | | | Group x Time | F (2,56) = 19.01 | <0.0001 |
| | c | Unpaired t-test | | t (28) = 6.54 | <0.0001 |
| | d | Two-way ANOVA (repeated measures) | Group | F (1,28) = 8.82 | 0.006 |
| | | | Time | F (2,56) = 10.55 | 0.0001 |
| | | | Group x Time | F (2,56) = 0.98 | 0.38 |
| | e | Mann-Whitney | | U = 53.50 | 0.027 |
| | f | Unpaired t-test | | t (28) = 5.33 | <0.0001 |
| | g | Unpaired t-test | | t (28) = 0.20 | 0.84 |
| | | Linear regression | | r = 0.27 | 0.15 |
| | h | Unpaired t-test | | t (28) = 0.44 | 0.66 |
| | | Linear regression | | r = 0.16 | 0.39 |
| | i | Mann-Whitney | | U = 48 | 0.014 |
| | | Linear regression | | r = 0.33 | 0.07 |
| | j | Unpaired t-test | | t (28) = 4.78 | <0.0001 |
| | | Linear regression | | r = 0.49 | 0.006 |

| Figure | Panel | Statistical test | Effect | Statistic | p-value |
|---|---|---|---|---|---|
| 3 | c | Three-way ANOVA (repeated measures) | Virus | F (1,68) = 0.0006 | 0.98 |
| | | | Group | F (1,68) = 0.041 | 0.84 |
| | | | Time | F (2,136) = 8.14 | 0.0011 |
| | | | Virus x Time | F (2,136) = 5.84 | 0.004 |
| | | | Virus x Group | F (1, 68) = 1.77 | 0.19 |
| | | | Group x Time | F (2,136) = 9.82 | 0.0001 |
| | | | Virus x Group x Time | F (2,136) = 12.84 | <0.0001 |
| | d | Two-way ANOVA (repeated measures) | Virus | F ( 1,68) = 13.37 | 0.0005 |
| | | | Group | F ( 1,68) = 25.83 | <0.0001 |
| | | | Virus x Group | F ( 1,68) = 23.77 | <0.0001 |
| | e | Three-way ANOVA (repeated measures) | Virus | F (1,68) = 0.01 | 0.92 |
| | | | Group | F (1,68) = 0.48 | 0.49 |
| | | | Time | F (2, 136) = 32.65 | <0.0001 |
| | | | Virus x Time | F (2,136) = 0.97 | 0.38 |
| | | | Virus x Group | F (1,68) = 12.80 | 0.0006 |
| | | | Group x Time | F (2,136) = 0.11 | 0.89 |
| | | | Virus x Group x Time | F (2,136) = 5.54 | 0.005 |
| | f | Two-way ANOVA (repeated measures) | Virus | F (1,68) = 0.42 | 0.52 |
| | | | Group | F (1,68) = 0.56 | 0.46 |
| | | | Virus x Group | F (1,68) = 14.24 | 0.0003 |
| | g | Two-way ANOVA (repeated measures) | Virus | F (1,68)=9.88 | 0.0025 |
| | | | Group | F (1,68) = 18.40 | <0.0001 |
| | | | Virus x Group | F (1,68) = 30.66 | <0.0001 |
| | h | Two-way ANOVA (repeated measures) | Virus | F (1,24) = 0.94 | 0.34 |
| | | | Group | F(1,24) = 0.38 | 0.54 |
| | | | Virus x Group | F (1,24) = 6.74 | 0.016 |
| | i | Two-way ANOVA (repeated measures) | Virus | F (1,24) = 10.57 | 0.0034 |
| | | | Group | F (1,24) = 6.77 | 0.016 |
| | | | Virus x Group | F (1,24) = 8.05 | 0.0091 |
| | k | Two-way ANOVA (repeated measures) | Virus | F (1,17) = 0.10 | 0.75 |
| | | | Time | F (2,34) = 17.23 | <0.0001 |
| | | | Virus x Time | F (2,34) = 4.26 | 0.022 |
| | l | Mann-Whitney | | U=33 | 0.34 |
| | m | Two-way ANOVA (repeated measures) | Virus | F (1,17) = 1.17 | 0.29 |
| | | | Time | F (2,34) = 19.44 | <0.0001 |
| | | | Virus x Time | F (2,34) = 0.17 | 0.84 |
| | n | Unpaired t-test | | t(17) = 1.09 | 0.29 |
